# Detection of *Plasmodium* sporozoites in *Anopheles coustani* s.l; a hindrance to malaria control strategies in highlands of western Kenya

**DOI:** 10.1101/2021.02.10.430589

**Authors:** Ayuya Stephen, Kitungulu Nicholas, Annette O. Busula, Mark Kilongosi Webale, Elizabeth Omukunda

**Affiliations:** Department of Biological Sciences, Masinde Muliro University of Science and Technology, Kakamega Kenya; Kaimosi Friends’ University College, a constitute college of Masinde muliro University of Science and Technology, Maragoli Kenya; School of Health Sciences, Kirinyaga University, Kutus, Kenya

## Abstract

Re-emerging of high malaria incidences in highlands of western Kenya pose a challenge to malaria eradication efforts. *Anopheles coustani* is a sub-Saharan mosquito species implicated in transmission of malaria in many parts of Africa as a secondary vector. It is a zoo-anthropophilic species that has been assumed to be of negligible importance. A cross sectional study was carried out in April to June, 2020 in Eluche location, Mumias East sub-County, Kakamega County, Kenya to establish the contribution of *Anopheles coustani* in malaria transmission. Pyrethrum spray collections (PSC) and Centers for Disease Control (CDC) and prevention light traps were used for sampling mosquitoes. Mosquitoes were collected from both indoors; between 0700h and 1100h using PSC and outdoors between 1800h and 0700h using CDC light traps. All mosquitoes were identified morphologically and female *Anopheles’* heads and thorax were analyzed further using Polymerase Chain Reaction (PCR) for *Plasmodium* sporozoite. A total of 188 female *Anopheles* mosquitoes were collected from both PSC and CDC light traps. This constituted of; 80(42.55%) *An. coustani*, 52(27.66%) *An. funestus*, 47(25.00%) *An. maculipulpis*, 8(4.26%) *An. arabiensis* and 1(0.53%) *An. gambiae*. Malaria sporozoite detection was done to all the *Anopheles* female mosquitoes but only two *An. coustani* tested positive for *Plasmodium falciparum*. In conclusion, *Anopheles coustani* plays a major role in outdoor malaria transmission in Mumias East Sub-County of Kakamega County in Western Kenya.

## Introduction

*Anopheles gambiae, An. arabiensis, An. funestus, An. nili* and *An. moucheti* are the Africa’s known primary malaria vectors (1,3). These vectors play a major role in malaria transmission and its sustenance (3–5). Secondary malaria vectors on the other hand include *An. coustani, An. pharoensis, An. ziemnni, An. rivolorum* and *An. maculipulpis* (3,5,6). Very little has been documented on vector competence of secondary vectors as compared to primary vectors(7–9). They have been considered of minimal or no importance in malaria transmission, which is not the case any more (10). In-depth knowledge on the ecology and both feeding and resting behavior of these vectors is crucial in malaria control strategies (11–13). To contribute to strategic malaria control program, it is vital to understand malaria transmission in terms of prevalence, incidence and mortality (14,15). Currently, long-lasting insecticidal nets (LLINs) and indoor residual spraying (IRS) are the base plans for vector elimination (7,16–18), which are however less effective against exophilic and exophagic vectors including *Anopheles coustani*.

Transmission of malaria has been attributed to vector presence, distribution, abundance, competence, capacity, (19,20) and of late shift in vector composition (21). Even though we have evidence of outdoor transmissions in some parts of sub-Saharan Africa, the available entomological scrutiny and monitoring tools focus on indoor mosquito populations which have changed in composition and feeding behavior (22). Vector abundance determines malaria transmission hence disease prevalence (19,23).

Insecticide resistance especially of pyrethroids that are used for vector control is on the rise (24– 26). Outdoor malaria transmission phenomenon has been reported (28,29). Recent studies have also documented shift in vector behavior from late night biting to early evening and morning biting (28,29). Increased use of LLINs, IRS and ITNs has seen vector behaviour change to zoophagic, exophagic and exophilic (21). Stable transmission of malaria in western Kenya highlands is supported by the biome, both micro and macro-climates. Equally, population pressure has seen changes in land use; valley bottom crop cultivation, deforestation and clearance of swamps (30,31).

In view of increasing concerns over residual malaria transmission in Mumias East, western Kenya, there is great need for vector behavior knowledge in terms of species composition, feeding behavior and possible contribution to indoor and outdoor transmission of malaria (22). This study was carried out in four randomly selected villages in Mumias East Sub-County to detect the presence of *Plasmodium falciparum* in field captured *Anopheles coustani* mosquito as well as establish its distribution in the region. Most vector control measures do not target *An. coustani* as it is an exophagic and zoophilic species (32). This study revealed that *Anopheles coustani is the main* vector in the study area. This therefore shows that there is need for outdoor vector control measures.

## Materials and Methods

### Study area and design

A cross sectional survey was carried out at Eluche Location in Mumias East Sub-County of Kakamega County in western Kenya; Approximately 25 KM South-west of Kakamega town, the County headquarters, located 00.34120 E and 034.54727 N with an average elevation of 1315 metres above sea level. The terrain of the area is fairly undulating with swampy valleys slopping to the west. The area receives adequate bimodal rainfall with a monthly mean of 222 mm (33). Agricultural activities in the area include sugarcane farming and mixed small scale farming (livestock; cattle, goat sheep, rabbit and even poultry: horticulture; maize, beans, kales, cabbages and indigenous vegetables).

### Household survey

A total of 220 houses were used in the survey. These houses were randomly selected based on the presence of aquatic habitats, house accessibility and characteristic; grass thatched, iron roofed, screened, unscreened, new or old. Distance between the houses with similar characteristics ranged from 50 to 100 meters apart. Written consent was issued for signing to the household head or representative after giving a go ahead during survey for sampling. Where consent was not granted, the next house was picked for sampling.

### Mosquito sampling and processing

Standard Centers for Disease Control and prevention, (CDC) light-traps were used to trap mosquitoes. Two light traps were placed per house, one indoor and another outdoor. Indoor traps were placed one and a half meters above the ground within sleeping areas for both animals and people between 19:00 and 06:00 hrs. Outdoor traps on the other hand were equally placed one and a half meters above the ground at areas secluded for animals to sleep for the same duration. Mosquitoes resting indoors were sampled using pyrethrum spray catch (PSC) method. This was done in the morning between 07:00h to 11:00h each day (34). Before spraying, all food stuff were covered. White sheets were spread on the floor and furniture in the houses. All openings to the houses including doors, windows and eaves were well sealed to avoid escape of mosquitoes. The roofs and walls were sprayed with 0.025% pyrethrum emulsifiable concentrate with 0.1% piperonyl butoxide in kerosene. All dead and unconscious mosquitoes from each house were collected from the sheets ten minutes after spraying, put in collection bottles (one per house), labelled, packed in a cool box and transported to Masinde Muliro University of Science and Technology, biotechnology laboratory where sorting was done and recorded. Each day’s anopheline collection was partially identified morphologically to species and sex level and kept in 1.5 ml Eppendorf tubes containing 70% isopropanol for further analysis. Malaria vector species complexes were identified by polymerase chain reaction using Scott *et al*., protocols(35).

### Data management and analysis

Data were entered in spread sheets and cleaned using Microsoft Excel, 2007. Statistical analyses were performed using SPSS version 23 and Graph pad Prism version 6 statistical software. One way Analysis of Variance (ANOVA) was used to obtain variation in mean densities between species and collections (indoor and outdoor). P value was set at 0.05 and 95% confidence interval.

## Results

### *Anopheles* species composition

A total of 188 *Anopheles* mosquitoes were collected using CDC light traps; Outdoor collections (n=127) and indoor collections (n=61). There was high abundance of *An. coustani [n=*80, (63.0%), *P<0*.*0001*] followed by *An. maculipalpis* [n=47, (37.0%), *P*<0.0001] in outdoor collections compared to indoor. There was high abundance of *An. funestus*, [n=52, (85.2%), *P*<0.0001] followed by *An. arabiensis*, [n=8, (13.1%), *P*<0.0001] in indoor collections in comparison to outdoor. However, there was insignificant abundance of *An. gambiae* [n=1, (1.6%) *P*=0.324] in indoor collection compared to outdoor collection (Table 1).

**Table 1.**
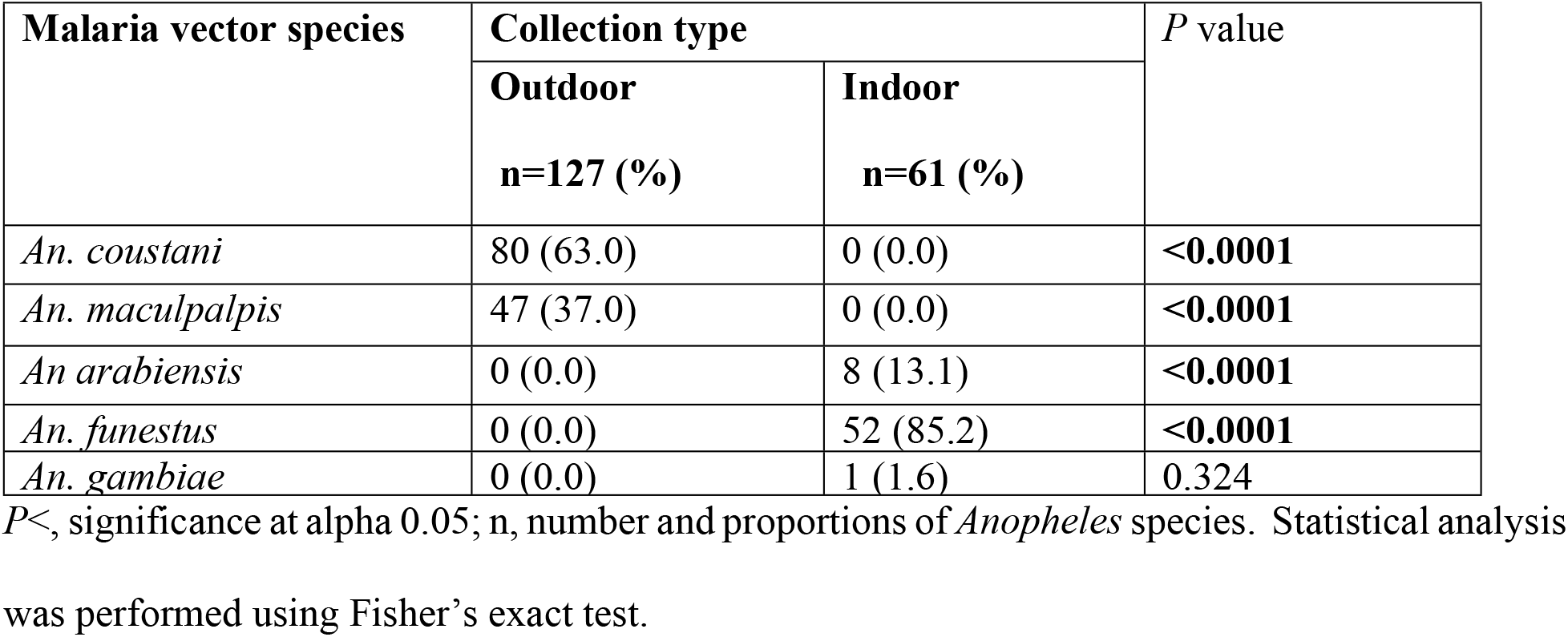
*Anopheles* species composition.

### *Palsmodium* sporozoites detection

All the 188 *Anopheline* mosquitoes captured were tested for presence of *Plasmodium* sporozoites. Only two *Anopheles coustani* tested positive with an infectivity rate of 2.5% (table 2). This result shows that *An. coustani* is majorly responsible for outdoor malaria transmission in this area. All the *An. coustani* used in this study were caught outdoors.

**Table 2.**
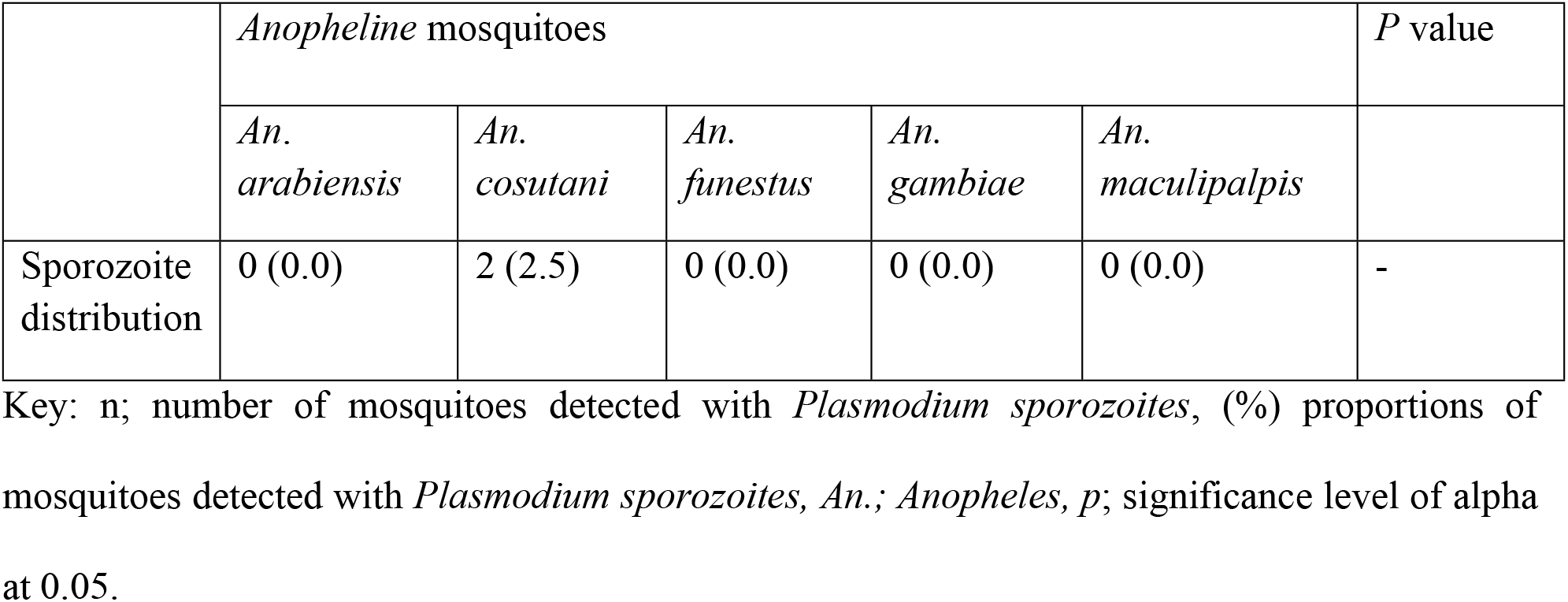
*Plasmodium* sporozoites detection analysis.

|  | *Anopheline* mosquitoes |  |  |  |  | P value |
| --- | --- | --- | --- | --- | --- | --- |
|  | *An. arabiensis* | *An. coustani* | *An. funestus* | *An. gambiae* | *An. maculipalpis* |  |
| Sporozoite distribution | 0 (0.0) | 2 (2.5) | 0 (0.0) | 0 (0.0) | 0 (0.0) | - |

Key: n; number of mosquitoes detected with *Plasmodium sporozoites*, (%) proportions of mosquitoes detected with *Plasmodium sporozoites*, *An.*; *Anopheles*, *p*; significance level of alpha at 0.05.

## Discussion

In sub-Saharan Africa, malaria infection remain the major cause of morbidity and mortality (36). Despite intensive studies on primary malaria vectors and the disease distribution, in-depth study on the role of secondary vectors on malaria transmission in the highlands of western Kenya is key (10). Much efforts have been concerted on the indoor vectors which primarily feed and rest indoors. Throughout the African continent, there is the use of LLINs and IRS which use domestic insecticides that effectively target endophagic and endophilic malaria vectors (10). Presently, most African countries do not have intrusions specifically targeting outdoor biting mosquitoes. Previous studies detected *Plasmodium* parasite in some secondary vectors across sub-Saharan Africa (3,4,10,37). *Anopheles coustani* is a species previously known to exhibit majorly zoophilic behavior (38). However, it was later established to play a role in malaria transmission due to its high anthropophilic tendencies (39). The implication of *An. coustani* in malaria transmission had only been studied in the coastal region of Kenya (10) where a total of five anopheline mosquito species; *An. arabiensis s*.*s, An. phunestus s*.*s, An. gambieae s*.*s, An. maculipulpis* and *An. coustani* were collected. Of these, the sub-Saharan Africa’s primary vectors: *An. funestus s*.*s, An. arabiensis s*.*s* and *An gambiae s*.*s* were caught indoors while the known primary vectors, *An. maculipulpis* and *An. coustani* were caught outdoors (21,40–43).

The present study found out that *An. coustani* had the highest proportion (42.55%) of the total anopheline female mosquitoes collected. This could be attributed to various factors that include; sugarcane plantations that influence the ecology of the vector through existence of numerous breeding sites. The study area is swampy but it is under reclamation through construction of drainage ditches. People live in clusters and stay outdoors up to very late in the evening in family social gatherings which enhance opportunistic feeding. The living standards of the people is low with grass thatched and semi-permanent houses that provide hiding places for the vectors. The very high abundance of the vector and detection of *Plasmodium* coupled with probable opportunistic feeding behavior enhances support of the role of the vector in malaria transmission (37). This study concurs with previous studies which have shown high composition of *Anopheles coustani* in outdoor collections (10,37). It however refutes some preceding studies where *An. coustani* were caught indoors while in this study, non was caught indoors (10,39,44,45). From the present study results, it is evident that Mumias East Sub-county of Kakamega County, Kenya experiences outdoor malaria transmission perpetuated by *An. coustani*.

This study was limited by the fact that it was a cross sectional study and consequently did not collect mosquitoes based on seasonality. It also did not look into vector species densities and distribution in relation to inoculation rates.

## Conclusion

The present study detected *Plasmodium* sporozoites in only *Anopheles coustani* mosquitoes. This means that *Anopheles coustani* plays an outstanding role in outdoor malaria transmission in this region. There is therefore need to evaluate the feeding patterns and social behavior of *Anopheles coustani* in highlands of Western Kenya for proper malaria vector management practices. Intervention programs should also include outdoor vector control strategies besides change of living routine.

## Data availability

The data used to support findings in this study are available from the corresponding author upon request.

## Conflicts of interest

The authors declare that there are no conflicts of interest.

## Acknowledgments

The authors would like to appreciate Mr, Maxwell GF. Machani of KEMRI Kisumu, Mr. Anthony Kibet of ICIPE Mbita for their technical support during the study

## Supporting information

SI Author list and contribution

S2 Suggested Reviewers

S3 Data

